# Knockdown of mitochondrial *atp1* mRNA by a custom-designed pentatricopeptide repeat protein alters F_1_F_o_ ATP synthase

**DOI:** 10.1101/2022.11.08.515711

**Authors:** Fei Yang, Lilian Vincis Pereira Sanglard, Chun-Pong Lee, Elke Ströher, Swati Singh, Glenda Guec Khim Oh, A. Harvey Millar, Ian Small, Catherine Colas des Francs-Small

**Author notes:** Author for correspondence: Catherine Colas des Francs-Small.

## Abstract

We show that a custom-designed RNA-binding protein binds and specifically induces cleavage of *atp1* RNA in mitochondria, significantly decreasing the abundance of the Atp1 protein and the assembled F_1_F_o_ ATP synthase in *Arabidopsis thaliana*. The transformed plants are characterized by delayed vegetative growth and reduced fertility. Five-fold depletion of Atp1 level was accompanied by a decrease in abundance of other ATP synthase subunits, lowered ATP synthesis rate of isolated mitochondria, but no change to mitochondrial electron transport chain complexes, adenylates or energy charge *in planta*. Transcripts for amino acid transport and a variety of stress response processes were differentially expressed in lines containing the PPR protein, indicating changes to achieve cellular homeostasis when ATP synthase was highly depleted. Leaves of ATP-synthase-depleted lines showed higher respiratory rates and elevated levels of most amino acids at night, most notably serine family amino acids. The results show the value of using custom-designed PPR proteins to influence expression of specific mitochondrial transcripts to carry out reverse genetics studies on mitochondrial gene functions and the consequences of ATP synthase depletion on cellular functions in *Arabidopsis*.

**One sentence Summary:** Knockdown of mitochondrial *atp1* mRNA by a custom-designed pentatricopeptide repeat protein alters F1Fo ATP synthase, plant growth and amino acid metabolism and ATP synthesis in *Arabidopsis thaliana*

## INTRODUCTION

Plant mitochondria are semi-autonomous organelles that produce adenosine triphosphate (ATP), the universal energy currency in the cell, through oxidative phosphorylation, whose final step is catalysed by the ATP synthase. Due to their endosymbiotic origins (Andersson et al., 2003), they contain their own genome, retaining about 65 functional genes. The scarcity of mitochondrial mutations, as well as the lack of reliable methods to transform mitochondria or knockdown expression of mitochondrial genes (Colas des Francs-Small et al., 2018; Kazama et al., 2019; Niazi et al., 2019) have made genetic analyses difficult (Kubo and Newton, 2008).

Although rare, spontaneous recombinations of the mitochondrial genome can generate chimaeric open reading frames (ORFs) encoding proteins that may cause pollen sterility (Chase, 2007; Mower et al., 2012).This phenomenon, known as cytoplasmic male sterility (CMS), is an important agronomic trait that has been widely used for plant hybrid seed production (Chen et al., 2017), and can be suppressed by nuclear restorer of fertility (*Rf*) genes (Hu et al., 2012; Huang et al., 2015). However, very few CMS genes have been functionally validated because of the lack of mitochondrial transformation strategies (Kazama et al., 2019), but mitochondrial ATP synthase subunit genes *atp1*, *atp4*, *atp6*, *atp8*, and *atp9* have often been found in CMS-associated loci (Hanson and Bentolila, 2004; Chen et al., 2017).

Within the large family of ATPases (Stewart et al., 2014), F_1_F_o_-ATP synthases in the inner mitochondrial membrane (complex V) produce ATP from adenosine diphosphate (ADP) and inorganic phosphate (Pi) by rotary catalysis, an essential process common to all forms of life (Kuhlbrandt, 2019). The complex consists of more than 17 different subunits (Supplemental Table 1) assembled into a soluble F_1_ sector and a membrane-embedded F_o_ sector, which are joined together by central and peripheral stalks (Artika, 2019). Proton translocation through the intermembrane space into the matrix drives the rotation of the F_o_ domain and the attached central stalk (Walker, 2013), and the conformational changes of the α (Atp1) and β (Atp2) subunits in F_1_ subsequently catalyse the synthesis of ATP (Srivastava et al., 2018). Although their bacterial and plastid counterparts are monomeric (Daum et al., 2010; Kuhlbrandt, 2019), mitochondrial ATP synthase complexes (~600 kD) arrange in rows of dimers, generating the characteristic curvature of the inner mitochondrial membrane known as cristae (Hahn et al., 2016; Gu et al., 2019), thus increasing the membrane surface and the density of respiratory complexes on a mitochondrial volume basis. ATP synthase dimers occupy the tips of the cristae whilst complex I and complex I-III super-complexes are limited to the flat sections of the mitochondrial inner membrane (Davies et al., 2011). Tetrameric structures have been reported in mitochondria from some mammals (Gu et al., 2019) and free-living ciliates (Flygaard et al., 2020) but not in plants.

To study energy metabolism related to CMS, several laboratories have attempted to knockdown nucleus-encoded subunits of ATP synthase. When induced during germination, antisense-RNA-mediated depletion of OSCP (ATP5) and γ (ATP3) subunits leads to seedling lethality, stressing the essential role of complex V. Lower levels of depletion resulted in altered leaf morphology, redox status, metabolism and gene expression (Robison et al., 2009). Depletion of the δ subunit by RNAi caused growth retardation, male sterility, female defects, decreased ATP synthase amounts, ROS accumulation and important metabolic changes (Geisler et al., 2012). Similar changes, as well as increased plant heat sensitivity, were observed in ATPd RNAi lines (Liu et al., 2021). Loss of ATP2 in *Chlamydomonas* altered mitochondrial and chloroplast ultrastructure and function (Lapaille et al., 2010). Mutations in *MGP1*, the gene encoding the F_A_d subunit of the mitochondrial ATP synthase lowered pollen viability (Li et al., 2010). Further studies have described the effects of expressing either an unedited copy of *atp9* (Busi et al., 2011) or fragments of the *atp4* transcript (Shaya et al., 2012), whilst others focused on mutants affected in the expression of mitochondrial ATP synthase subunit genes, such as *otp87* (Hammani et al., 2011; Colas des Francs-Small and Small, 2013).

In recent years, in the absence of a reliable mitochondrial transformation method, several indirect approaches aiming to alter mitochondrial gene expression have been attempted. Targeted knockdown of mitochondrial gene expression was achieved via tRNA-like ribozymes (Val et al., 2011; Sultan et al., 2016; Niazi et al., 2019). In another approach, CMS-associated genes *orf79* in rice and *orf125* in rapeseed were knocked out by using transcription activator-like effector nucleases (TALENs) targeted to mitochondria (Kazama et al., 2019). This method was also used to generate deletions in both copies of the *atp6* gene (Arimura et al., 2020).

We developed a different approach for reverse genetics in plant mitochondria, which successfully induced cleavage of *nad6* transcripts by a custom-designed pentatricopeptide repeat (PPR) protein (Colas des Francs-Small et al., 2018). PPR proteins are organelle RNA-binding proteins that have more than 400 members in most species of land plants (Barkan and Small, 2014). The protein family is divided into subgroups according to the length and disposition of their repeated motifs (Cheng et al., 2016). They bind organelle transcripts in a sequence-specific manner (Yin et al., 2013; Yan et al., 2019) and affect the editing, processing, splicing or translation of the target RNA (Schmitz-Linneweber and Small, 2008). Most restorer of fertility (*RF*) genes encode PPR proteins (Kim and Zhang, 2018) and form a small clade together with restorer-of-fertility-like (RFL) proteins (Fujii et al., 2011; Melonek et al., 2016; Anisimova et al., 2019; Melonek et al., 2019). RF proteins interact with and block translation of CMS transcripts by inducing cleavage, preventing ribosome translocation (Dahan and Mireau, 2013; Gaborieau et al., 2016; Wang et al., 2021)]. Among 26 RFL proteins in *Arabidopsis thaliana*, several functionally characterised members are also known to induce cleavage of mitochondrial transcripts (Jonietz et al., 2010; Holzle et al., 2011; Arnal et al., 2014; Stoll et al., 2017).

The PPR code describing how two amino acids in each PPR protein repeat recognise each base of its target RNA provides a strategy to design PPR proteins for plant mitochondrial RNA manipulation (Barkan et al., 2012; Yan et al., 2019). We previously redesigned the RFL protein RPF2 (also known as RFL6) to bind within the *nad6* transcript coding sequence, promoting its cleavage (Colas des Francs-Small et al., 2018). This led to undetectable amounts of Nad6 subunit and consequently undetectable amounts of assembled complex I, the first complex of the respiratory chain. The high specificity of PPR proteins for their targets allowed us to effectively and precisely knockdown *nad6*, as shown by the few off-target effects. If generalisable, this method would be very useful for altering mitochondrial transcript abundance and studying the control of expression of the building blocks of multi-subunit enzyme complexes and their assembly process.

In this work, we show that a modified RFL protein (RPF2-*atp1)* designed to bind the coding sequence of mitochondrial *atp1* can effectively knockdown *atp1* transcripts *in vivo*, causing delayed growth and reduced fertility in *Arabidopsis*. How these plants can develop and reproduce under these deleterious conditions is explored.

## RESULTS

### RPF2-*atp1* Protein Design and Primary Transformant Screening in T1 generation

The native RPF2 protein (At1g62670) is composed of 16 PPR motifs (Figure 1A). It participates in mitochondrial mRNA maturation by binding the 5′-untranslated regions (UTRs) of *cox3* and *nad9* and inducing cleavage (Jonietz et al., 2010). Using the online program EMBOSS: fuzznuc, we found a sequence at position + 1330-1346 in the coding sequence of *atp1* (coordinates 67,292-67,277 on the Col-0 mitochondrial genome BK010421 (Sloan et al., 2018)) with only 5 differences to the predicted RPF2-binding site in the *nad9* transcript (Figure 1A). The coding sequence of RPF2 was modified to produce a designed protein (RPF2-*atp1*) able to recognize the *atp1* mRNA target sequence (Figure 1A and Supplemental Figure 1). *A. thaliana* Col-0 plants were transformed with the synthetic construct via *Agrobacterium* infection. Integration of the construct into the genome was checked in primary transformants (T1) by genomic PCR (Supplemental Figure 2A). Fourteen out of 32 independent transformants carrying the RPF2-*atp1* construct displayed slow growth and delayed flowering (Figure 1B and 1C). Plants carrying the native RPF2 cDNA (Colas des Francs-Small et al., 2018) were used as controls. The expression of the RPF2-*atp1* construct in transgenic lines was verified by RT-PCR and western blotting (Supplemental Figure 2B and 2C). An RT-qPCR experiment performed on four independent RPF2-*atp1* lines (RPF2-*atp1-*2, 9, 16 and 19) and on the control plants confirmed that high expression of RPF2-*atp1* reduced *atp1* transcript accumulation (Supplemental Figure 2B).

**Figure 1.**
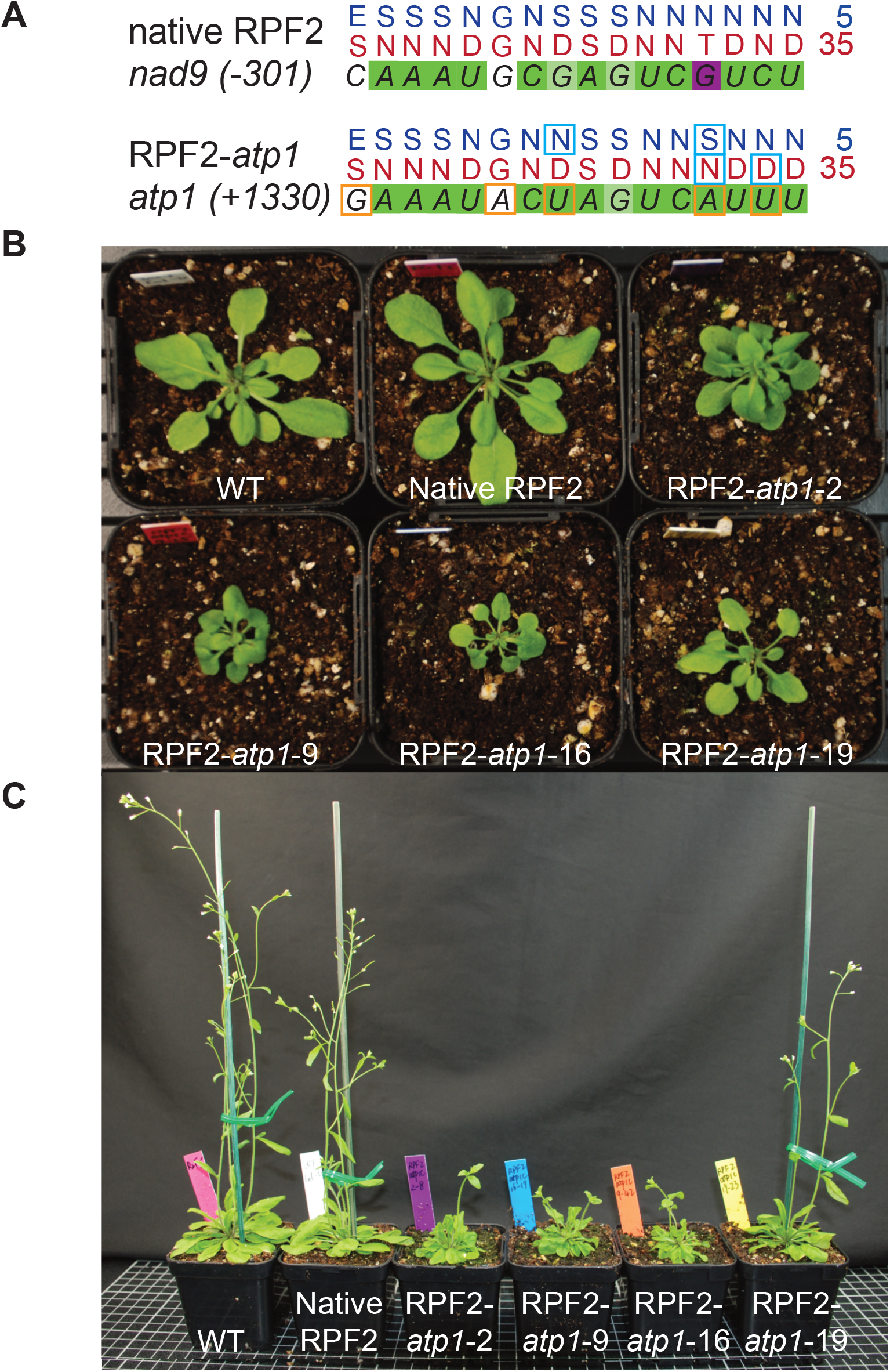
Targeting the *atp1* Transcript with a Modified RPF2 Protein Leads to Delayed Plant Growth and Curled Leaf Phenotype. (**A**) RNA targets for RPF2 (*nad9* 5’ UTR) and RPF2-*atp1* (*atp1* CDS) and respective binding predictions. Dark green squares represent a perfect match, light green a partial match, white a neutral match and magenta a mismatch according to the PPR code. The differences between RNA targets are highlighted by orange squares and the modifications in the protein by cyan squares. (**B**) and (**C**) show the phenotypes of four lines (T2 generation) transformed with the RPF2-*atp1* and native RPF2 as compared with WT. (**B**) 4-week-old rosettes; (**C**) 6-week-old plants grown under 18 h photoperiod.

### The *atp1* Transcript is Cleaved in RPF2-*atp1* Plants

To check for cleavage of the *atp1* transcript, northern blotting was carried out in the T1 generation using *atp1*-specific biotinylated oligonucleotide (Figure 2A). Probe 404 AS (upstream of the predicted binding site) hybridized to a single ~2044 nt transcript in WT samples corresponding to the expected size of the mature *atp1* transcript (1524 nt of coding sequence and around 520 nt of 5’- and 3′-UTRs), but in most RPF2-*atp1* plants, this hybridization signal was very weak and accompanied by an additional faint signal migrating around 1760 nt (asterisks in Figure 2A), suggesting that the *atp1* transcript was indeed cleaved in the RPF2-*atp1* plants. With probe 1457AS (hybridizing 3′ of the predicted binding site), a very strong signal corresponding to a potential cleavage product was detected only in the RPF2-*atp1* lines, migrating at ~280 nt in the RPF2-*atp1*-2, RPF2-*atp1*-9 and RPF2-*atp1*-16 plants (Figure 2A). In RPF2-*atp1*-19 plants, two bands of ~2044 nt and ~280 nt were detected, indicating that *atp1* transcripts were partially cleaved. Circular RT-PCR (cRT-PCR) experiments were conducted in two independent lines (RPF2-*atp1*-9 and RPF2-*atp1*-16), to precisely locate the cleavage site. The cRT-PCR products were cloned and sequenced. Multiple transcript ends could be mapped within a region encompassing 55 nucleotides beginning at or near the end of the predicted RPF2-*atp1* binding site (Figure 2B).

**Figure 2.**
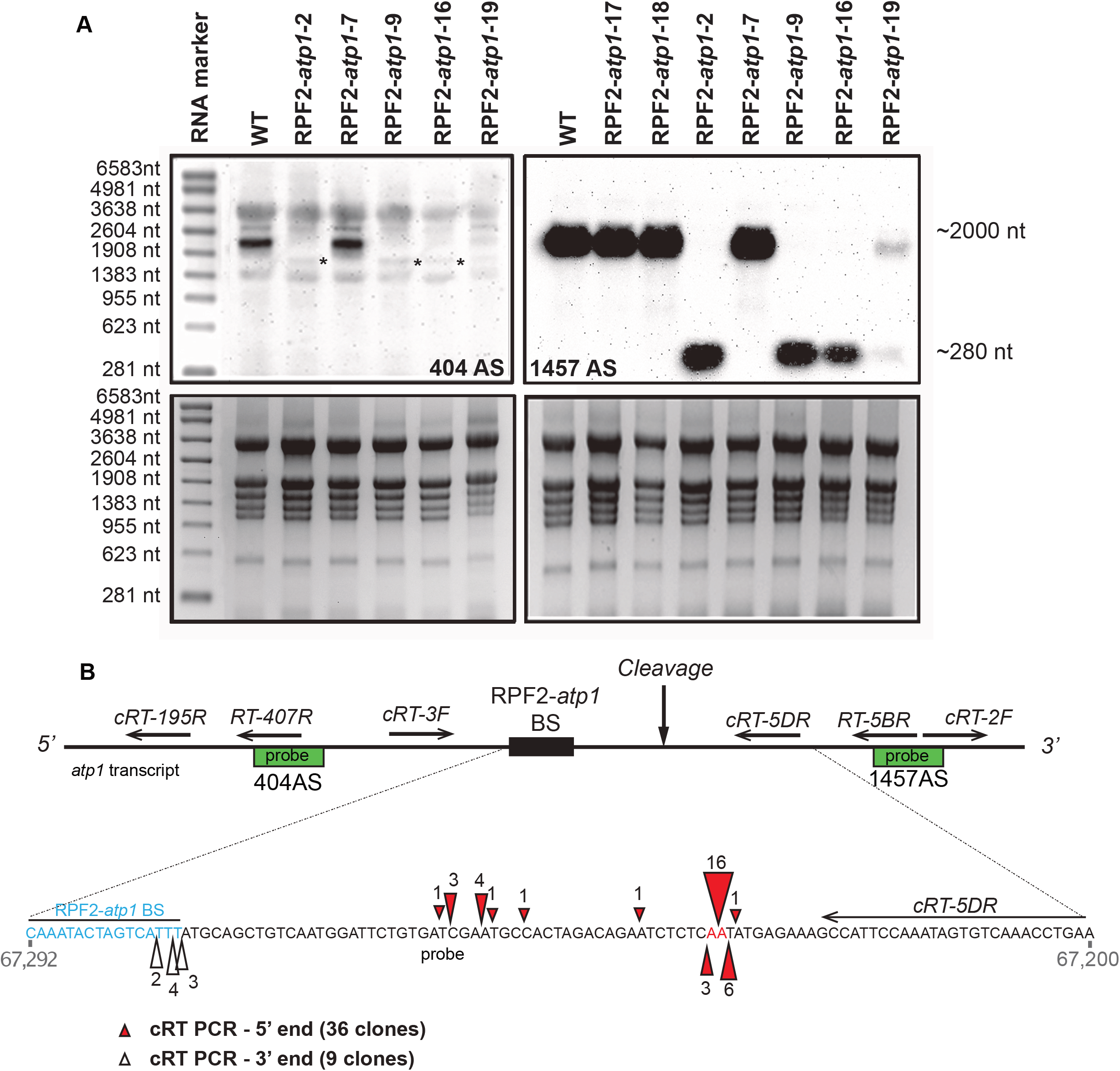
The *atp1* Transcript is Cleaved in the Plants Transformed with the RPF2-*atp1* Constructs. (**A**) Northern blots of leaf RNA isolated from several transformants and WT plants hybridised with *atp1* probes 404AS (top left) and 1457AS (top right), respectively located upstream and downstream of the RPF2-*atp1* predicted binding site (BS). The bottom panels show the gels stained with Ethidium bromide. The molecular weight marker sizes are indicated on the left-hand side. The asterisks on the 404 AS blot show the faint cleavage bands around1760 nt. (**B**) Location of *atp1* cleavage in the RPF2-*atp1* plants from cRT-PCR results. The top panel indicates the position of the northern probes used in (**A**) (green boxes) and cRT PCR primers relative to the predicted binding site (not to scale). The coordinates of the enlarged region on the Col-0 mitochondrial genome BK010421 are 67,292 to 67,200 (reverse strand).The RPF2-*atp1* binding site (67,277-67,292) and the main cleavage site are highlighted. Red triangles indicate the 5’ ends of the cleaved products and white triangles the 3’ ends of the cleaved products as determined by cRT-PCR. The ends of the 36 clones aligned suggest cleavage between bases 67,224-67,225. The figures near the triangles indicate the numbers of clones obtained.

### RPF2-*atp1* Plants Have Decreased Abundance of Complex V and Atp1

Blue native polyacrylamide gel electrophoresis (BN-PAGE) was performed on crude leaf membranes from T2 plants of the transformants to check the integrity of the F_1_F_o_ ATP Synthase complex (Figure 3A). Western blotting of the BN-PAGE gel and probing with an antibody raised against Atp1 revealed assembled complex V in the mitochondria of WT plants, WT plants transformed with native RPF2 (referred to hereafter as native RPF2 plants) and RPF2-*atp1*-19-1 and 19-2 plants, but reduced amounts in mitochondria from RPF2-*atp1*-2 plants and almost undetectable levels in mitochondria from RPF2-*atp1*-9, RPF2-*atp1*-16 and *otp87* plants (Figure 3A). Further analysis of respiratory complex subunits by SDS-PAGE and western blotting showed that Nad9 (complex I), RISP (complex III), and Cox2 (complex IV) subunits were unchanged in abundance in RPF2-*atp1* mitochondria, but Atp1 was undetected in mitochondria of RPF2-*atp1*-9 and RPF2-*atp1*-16 plants using this technique (Figure 3B) and lower in abundance in RPF2-*atp1*-2 and RPF2-*atp1*-19 mitochondria than in WT mitochondria. Because the *atp1* transcript cleavage is partial in the RPF2-*atp1*-19 line (Figure 2), a small amount of Atp1 was detected, allowing assembly of some complex V (Figure 3A). Furthermore, alternative oxidase (AOX) and HSP70 were more abundant in RPF2-*atp1*-9 and 16 plants (Figure 3B), as observed for other mutants with altered complex V function (Hammani et al., 2011; Kerbler et al., 2019).

**Figure 3.**
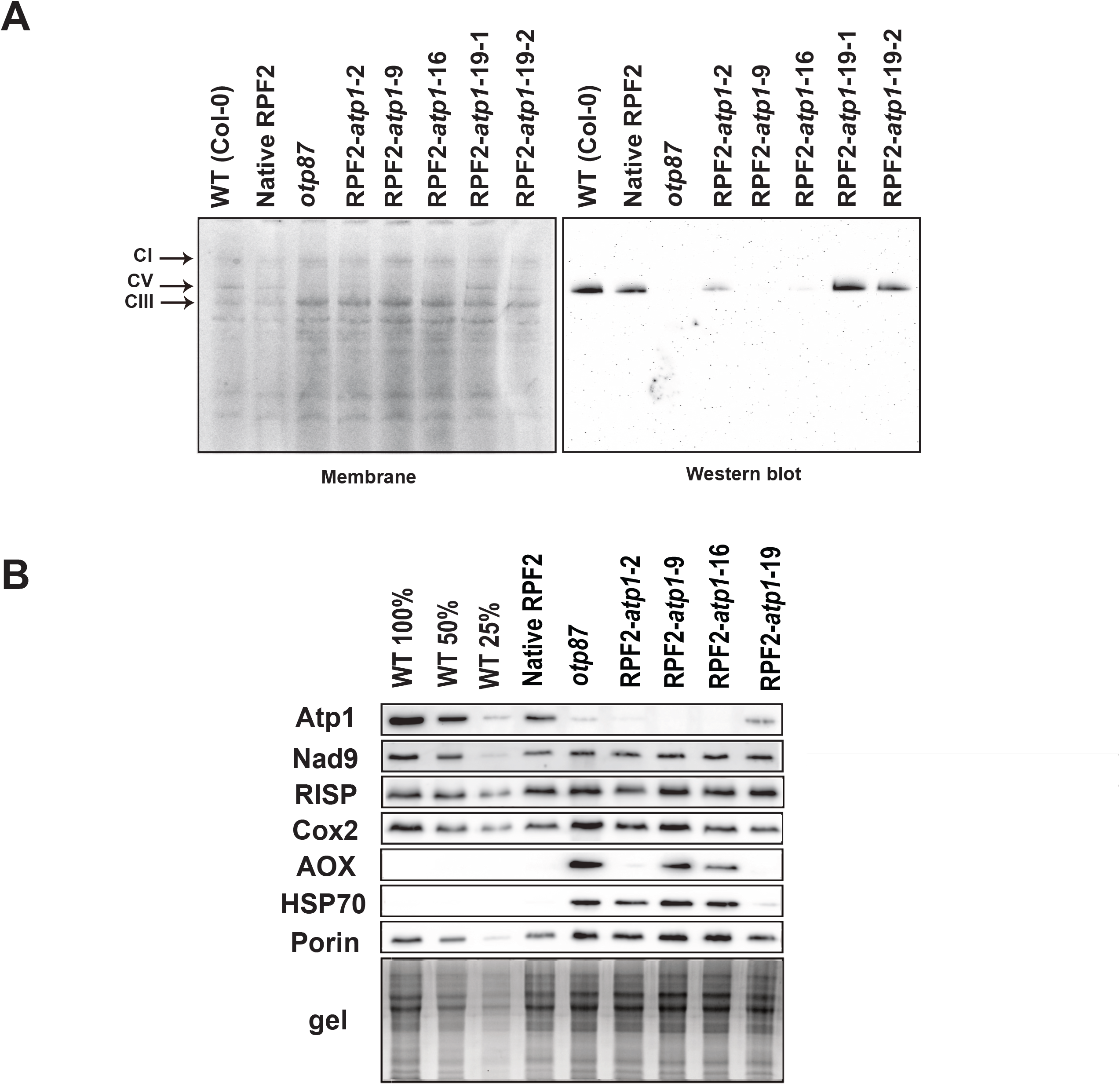
RPF2-*atp1* Plants Lack Atp1 Subunit and Assembled Respiratory Complex V. (**A**) Separation of crude membrane protein complexes by Blue Native PAGE of four RPF2-*atp1* transformants as compared with WT and *otp87* (an *atp1* editing mutant). The left panel shows the stained membrane after transfer and the right panel the western blot probed with an anti-Atp1 antibody. The black arrows show complex I (CI), V (CV) and III (CIII). (**B**) Western blots of mitochondrial proteins of four RPF2-*atp1* transformants as compared with WT, native RPF2 and *otp87* separated by SDS PAGE.

### The Modified Plants Show Small Rosettes, Delayed Growth, Reduced Fertility but Increased Respiration Rates

In T3 generation, the traits of the modified plants were comprehensively investigated. In week 4, the rosette diameters in RPF2-*atp1*-2, 9, 16 and 19 were only half those of WT and native RPF2 plants (Figure 1B, Supplemental Figures 3 and 4). In week 6, RPF2-*atp1*-2, 9 and 16 still had much smaller rosettes, shorter plant height than control plants, RPF2-*atp1*-19 was similar to WT and native RPF2 plants, in accordance with the *atp1* transcript partial cleavage. Leaf numbers per plant were similar in all genotypes in week 4 but stopped increasing in the modified plants in week 6. The rosettes of RPF2-*atp1*-2, 9 and 16 developed downwards curled leaves, clearly different from RPF2-*atp1*-19 and control plants (Supplemental Figure 3D). Bolting and flowering were delayed in the transgenic lines. RPF2-*atp1*-9 plants were most affected, with few very short siliques, resulting in low seed yield per plant (Supplemental Figures 3B, 3C and 4A). RPF2-*atp1*-2 plants showed some instability across generations and were not analysed further. Root lengths of 10-day-old seedlings grown on vertical plates (Supplemental Figures 3E and 4B) were significantly shorter in RPF2-*atp1* lines than in control (*p* = 7.5 10^−44^ for RPF2-*atp1*-16; *p* = 1.1 10^−29^ for RPF2-*atp1*-9).

The oxygen consumption rates of leaves from T3 plants (16/8 hours, day/night) for 6 weeks were measured by a fluorophore-based oxygen sensor. The average oxygen consumption rates of RPF2-*atp1*-9 and RPF2-*atp1*-16 plants were significantly higher than those of WT and native RPF2 plants during the 12 hours of measuring time, whilst RPF2-*atp1*-19 average respiration rates were close to the controls (Supplemental Figure 5A). To minimize developmental stage differences between genotypes, short day conditions (8/16 hours, day/night) were applied, and molar O_2_ consumption was calculated for RPF2-*atp1*-9 and −16 (34 and 47 plants respectively), the *otp87* mutant (18 plants) (Hammani et al., 2011) and the native RPF2 and WT controls (31 and 28 plants respectively). The RPF2-*atp1*-9 and −16 lines displayed 24% higher oxygen consumption rate than the controls, slightly less than *otp87* (29.7%), but significantly higher rates than the controls (Supplemental Figure 5B).

### RNA-Seq Confirms Reproductive Development Defects in the RPF2-*atp1*-9 Plants and Highlights Their High Level of Stress

RNA-seq data obtained from 3 biological repeats of the RPF2-*atp1-*9 line (S9 1-3) compared to WT (C2-4) were analysed in the T4 generation. Out of 18643 transcripts, 5683 were found to be differentially accumulated using a false discovery rate threshold of 0.1, of which 2831 were more abundant in *atp1* and 2852 were more abundant in WT (Supplemental Table 2). A set of 11 of the GO terms most significantly over-represented in the annotations of genes whose expression was lower in RPF2-*atp1*-9 plants are associated with reproductive development (Figure 4A, Supplemental Table 3). Forty-three out of 45 genes implicated in pollen exine formation, 32 out of 62 genes implicated in pollen tube growth and 27 out of 48 implicated in stamen development were significantly less expressed in the RPF2-*atp1*-9 plants than in WT, in agreement with their reduced fertility (Supplemental Figures 3B, 3C and 4A). GO terms relating to plant growth and development were also highly represented (cell wall organization, post-embryonic development, petal development, meristem development, cell cycle, cell fate, organelle organization) as well as protein metabolism and lipid transport. Most of the 29 GO terms significantly over-represented in the terms annotating genes more expressed in RPF2-*atp1*-9 plants (Figure 4B, Supplemental Table 4) were related to biotic and abiotic stress responses, consistent with the increased levels of AOX and HSP70 observed by western blotting (Figure 3B). Interestingly, processes such as protein targeting to membrane and response to unfolded protein were also over-represented as well as amino acid transport.

**Figure 4.**
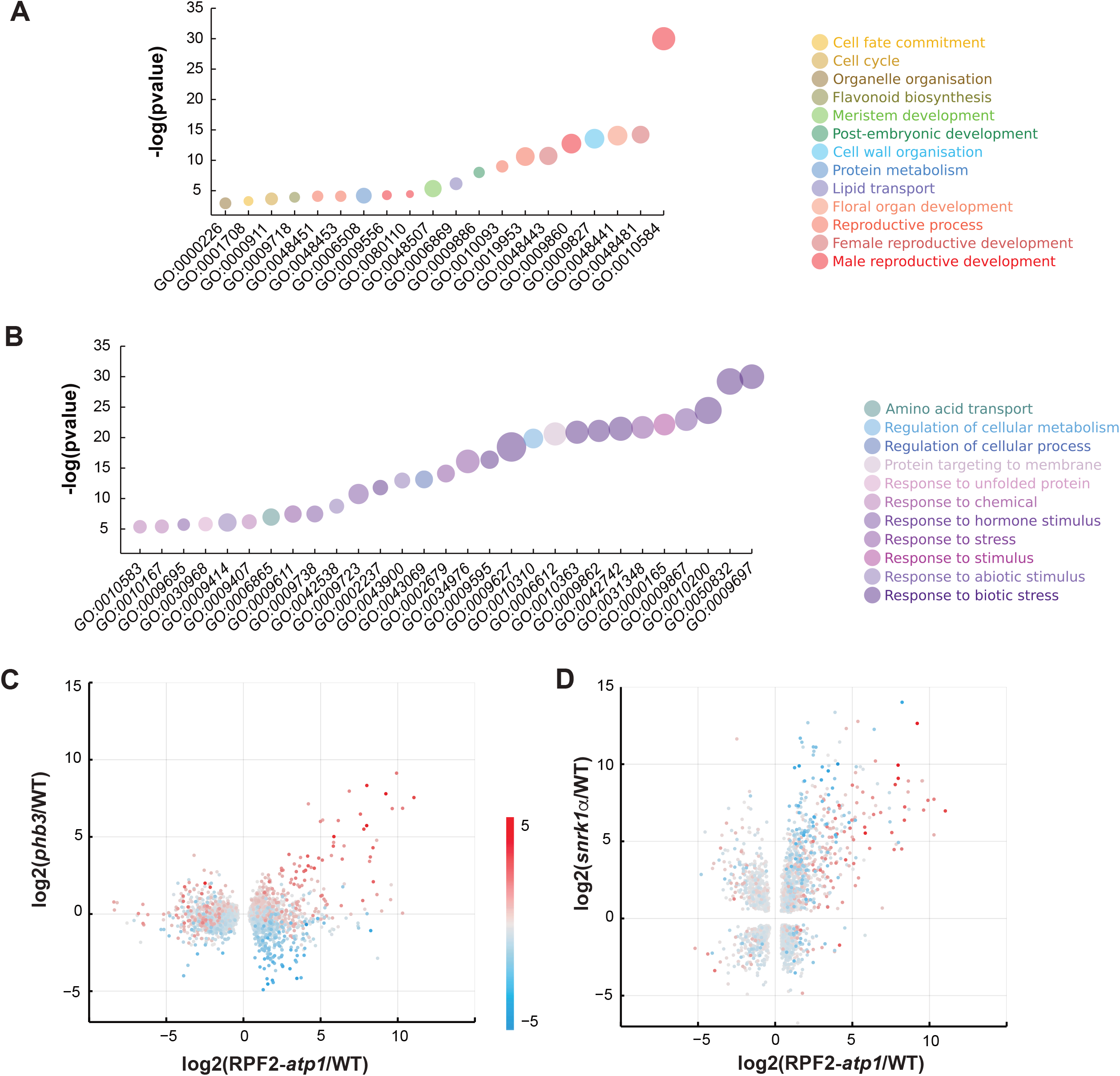
RNA-seq analysis of Wild-Type and RPF2-*atp1* Plants. (**A, B**) GO Biological Process terms most significantly associated with genes more expressed in WT (**A**) or in RPF2-*atp1* plants (**B**).Colours indicate groupings of GO terms into broader categories. The size of the markers is proportional to the number of differentially expressed genes in each category. The y axis indicates the degree of statistical significance, higher values being more significant. (**C**) A scatter plot comparing the transcript differences in RPF2-*atp1* and *phb3* mutants (x-axis is log_2_ fold-change between RPF2-*atp1* and WT, y-axis is log_2_ fold-change between *phb3* and WT). Only transcripts that are significantly differentially expressed in RPF2-*atp1* mutants are included. The marker colour represents the effect of the ANAC017 transcription factor (calculated as the log_2_ ratio in expression between *phb3* mutants and *phb3 anac017* double mutants). (**D**) A scatter plot comparing the transcript differences in RPF2-*atp1* and *snrk1α* mutants (x-axis is log_2_ fold-change between RPF2-*atp1* and WT, y-axis is log_2_ fold-change between *snrk1α* and WT). Only transcripts that are significantly differentially expressed in both mutants are included. The marker colour represents the effect of the ANAC017 transcription factor (calculated as the log_2_ ratio in expression between *phb3* mutants and *phb3 anac017* double mutants).

### RNA-seq shows involvement of the ANAC017 transcription factor and SNRK1 kinase in RPF2-*atp1* line gene expression changes

To assess if transcript increases observed in RPF2-*atp1* lines could be related to the well-documented mitochondrial retrograde signals mediated by the transcription factor ANAC017 (De Clercq et al., 2013; Ng et al., 2013), we plotted our RPF2-*atp1* data against RNA-seq data obtained from *phb3* and *phb3 anac017* double mutants (Van Aken et al., 2016). Most of the transcripts strongly induced in RPF2-*atp1* are also induced in *phb3* and are ANAC017-dependent, but a second group of transcripts induced in RPF2-*atp1* are not induced in *phb3* and not dependent on ANAC017 (Figure 4C). In attempting to understand what mediates the induction of these transcripts in the RPF2-*atp1* plants, we compared our RNA-seq data to similar data from a *snrk1α* mutant deficient in a SNRK1 kinase involved in metabolic adaptation to low energy supply (Pedrotti et al., 2018). As for the comparison with *phb3*, ANAC017-dependent transcripts are induced in both RPF2-*atp1* and *snrk1α*, but more so in RPF2-*atp1* (Figure 4D). Interestingly, the ANAC017-independent group of transcripts induced in RPF2-*atp1* plants but not in *phb3* are even more strongly induced in *snrk1α*.

### Analysis of Mitochondrial Transcripts Did Not Reveal Any Significant Off-Target Effects of the RPF2-*atp1* Protein

The RNA-seq data was also mined to look for off-target effects due to potential binding of RPF2-*atp1* on transcripts other than the targeted *atp1*. Confirming the RT-qPCR results, the *atp1* transcript was significantly less abundant in RPF2-*atp1* samples (about four-fold lower; p = 0.004) (Supplemental Figure 6). The RNA-seq data was also analysed to specifically search for cleavage events generated by the RPF2-*atp1* protein. Figure 5A shows the relative frequency of RNA-seq 5’ ends across the whole mitochondrial transcriptome for WT and RPF2-*atp1*-9 samples; the profiles for both genotypes are very similar overall. Only two prominent peaks are significantly higher in the RPF2-*atp1-9* samples (Figure 5B, C), one in the *atp1* transcript at position 67233 (reverse strand), corresponding to the cleavage site induced by the RPF2-*atp1* protein (as determined by cRT-PCR, Figure 2B), and the other in *nad2* intron 2 (position 98564, reverse strand). This second peak does not represent an RNA cleavage site, but rather the 5’end of the half-intron (as *nad2* intron 2 is trans-spliced). The increase in abundance of this peak suggested that the splicing efficiency of *nad2* intron 2 might be decreased in RPF2-*atp1-9* plants, so we calculated the splicing efficiencies for all intron-containing mitochondrial transcripts in both genotypes (Supplemental Figure 7). The splicing of almost all introns was found to be significantly lower in RPF2-*atp1*-9 plants than in WT. Apparent relative editing rates in WT as compared to RPF2-*atp1*-9 were also calculated for 461 mitochondrial editing sites within coding sequences and found to be generally lower in the modified plants (Supplemental Figure 8).

**Figure 5.**
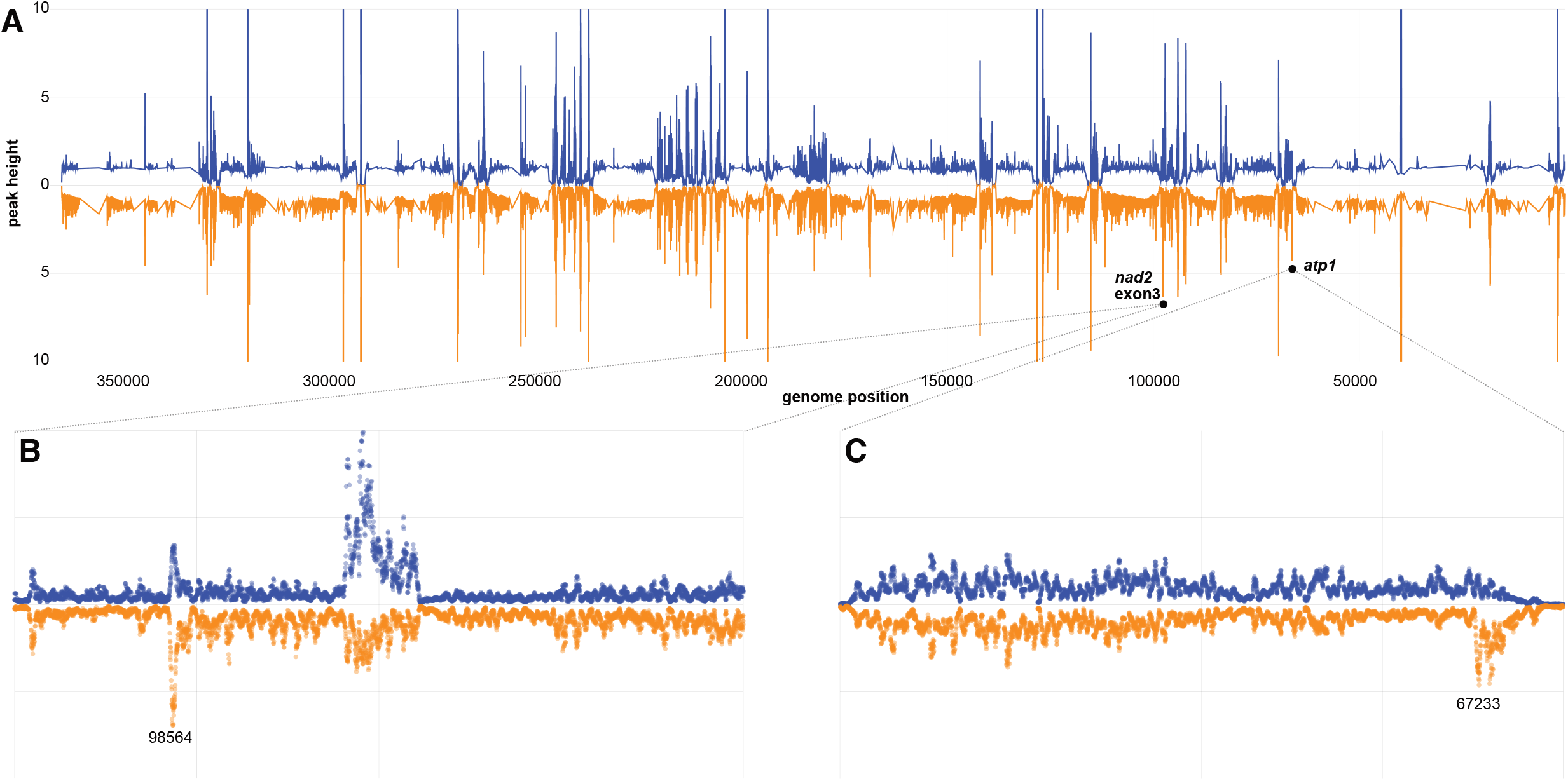
Comparative Mapping of RNA-Seq 5’ Ends Confirms Cleavage of *atp1* Transcripts Without Significant Off-Target Cleavages. (**A**) Relative frequency of RNA 5’ ends mapping to the reverse strand of BK010421 in WT (blue) and RPF2-*atp1* (orange) plants. The data has been smoothed and normalized relative to the local background, is a mean of 3 samples for each genotype, and is shown from 5’ to 3’ (BK010421 coordinates 367808–1). The RPF2-*atp1* data is plotted as a mirror image to facilitate comparison to the WT pattern. (**B**) Plot of the same data in the region (99000–97000) but now showing individual data points for each sample at single-nucleotide resolution. The peaks in the centre of the panel indicate the extent of *nad2* exon 3 (BK010421 coordinates 95238–94666). The peak in RPF2-*atp1* samples at 98564 (*p* = 0.019) is one of only 2 peaks in the mitochondrial transcriptome to be significantly more prominent in RPF2-*atp1* than in WT. We believe it to indicate the 5’ end of unspliced *nad2* exon3 transcripts. (**C**) Plot of the data in the region (69000– 67000) showing individual data points for each sample at single-nucleotide resolution. This region covers the *atp1* gene (BK010421 coordinates 68621–67098). The peak in RPF2-*atp1* samples at 67233 is significantly more prominent in RPF2-*atp1* than in WT (*p* = 0.0099). We believe it to indicate the cleavage site(s) induced by RPF2-*atp1*.

### Proteomics and Metabolomics Analyses

Figure 6A describes the assembly pathway of ATP synthase (adapted from (Röhricht et al., 2021)) and highlights the other subunits that make up ATP synthase in plant mitochondria. A quantitative untargeted mass spectrometry approach was used to explore potential changes in mitochondria caused by decrease in the ATP synthase complex and to check levels of the subunits of the respiratory complexes in RPF2-*atp1-9* and RPF2-*atp1-16* as compared with WT and native RPF2. Out of 410 mitochondrial proteins in our analysis, the most impacted in abundance were Atp1 (target of RPF2-*atp1*) and most other components of the ATP synthase complex (Figure 6B, Supplemental Table 1, Supplemental Figure 9). All components of the F_1_ domain (subunits α, β, γ, δ and ε) were decreased to 15-25% of WT, so we can confidently conclude that F_1_ assembly was greatly affected. Most subunits of the peripheral stalk (subunits b, d, OSCP, f, F_A_d and i/j) followed the same trend, suggesting that the F_o_ domain was not able to assemble properly either. Unfortunately, we could not detect with mass spectrometry the subunits a (Atp6), 8 (Atp8) and c (Atp9), all highly hydrophobic components of the F_o_ domain.

**Figure 6.**
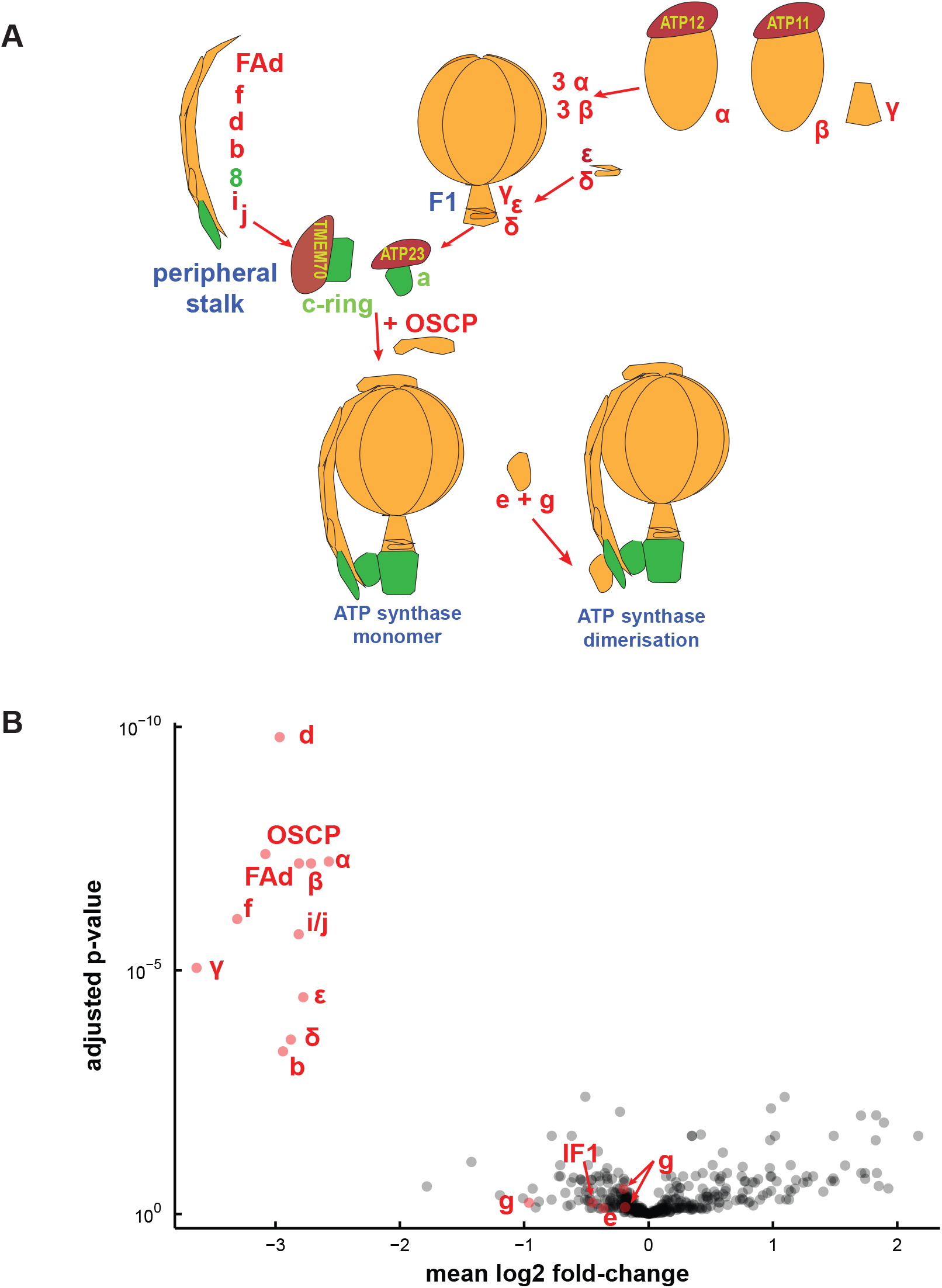
Most Subunits of the ATP Synthase Complex Are Less Abundant in RPF2-*atp1* Transformants. (**A**) Simplified representation of the ATP synthase assembly pathway. The subunits found in our proteomics study are labelled in red, the others are labelled in green. (**B**) A ‘volcano’ plot of relative protein abundances as estimated by quantitative untargeted mass spectrometry (MS) from samples of WT and RPF2-*atp1* crude mitochondrial pellets. The x axis indicates mean log_2_ fold-difference (WT/ RPF2-*atp1*), the y axis indicates the p-value for the hypothesis of equal relative abundance, i. e. points towards the top-left of the plot are significantly less abundant in RPF2-*atp1* samples. The plot shows data for 410 proteins of which 17 (in red) are associated with the ATP synthase complex. The plot is based on MS data from 8 RPF2-*atp1* samples and 8 phenotypically WT samples (4 from WT plants, 4 from plants expressing the native RPF2).

Unexpectedly, three ATP-synthase-associated proteins did not vary: g (ATP21), e (ATP20), both involved in the dimerization of the F_o_ sub-complex (Zancani et al., 2020; Röhricht et al., 2021) and IF1, an inhibitor of ATP synthase activity (Chen et al., 2020). No significant protein changes were detected in subunits of electron transfer chain complexes (Figure 6B; Supplemental Figure 9), in accordance with the western blot results (Figure 3B). Analysis of the RNA-seq data showed induction of transcripts for two non-phosphorylating bypasses of the electron transport chain, namely alternative oxidase (*AOX1a* and *AOX1d*) and rotenone insensitive NADH dehydrogenase (*NDA1*, *NDB2*, *NDB4*).

In order to understand how, despite only having 15-25% of the mitochondrial ATP synthase found in WT, most RPF2-*atp1* transformants can grow, flower and set seed, we measured total adenylates in roots at the end of the dark period. Root tissue was chosen to minimize the fraction of adenylates produced by the chloroplast ATP synthase. These measurements (5 biological repeats) showed that the total root abundance of ATP, ADP and AMP in the transformants were surprisingly close to those in the control plants with an ATP/ADP ratio of approximately 1.8 and an adenylate charge ((ATP+0.5*ADP)/(ATP+ADP+AMP)) of 0.8 (Supplemental Figure 10A, Supplemental Table 5). ATP synthesis rates were directly evaluated in purified substrate-energized mitochondria and found to be 44% and 57% slower in RPF2-*atp1*-16 and RPF2-*atp1*-9 respectively than in native RPF2 (WT) seedlings (Supplemental Figure 10B). This shows that while ATP synthase abundance is substantially lowered, it only has a moderate metabolic control coefficient (~0.4) on ATP synthesis rate, potentially explained the ability of RPF2-*atp1-9* and *-16* to grow at a slower rate and maintain adenylate pools *in vivo*.

To gain further insight into the apparent changes in metabolic processes, and changes in amino acid transport processes indicated by transcript profiling (Figure 4B), absolute organic acid and amino acid levels were measured by LC-MS in seedlings at the end of a dark period (Supplemental Figure 11). This revealed significant changes in metabolite abundances between the RPF2-*atp1* and WT plants. Citrate and malate accumulated but fumarate was less abundant in the RPF2-*atp1* plants compared to native RPF2 (WT) controls. All amino acids except threonine accumulated more in RPF2-*atp1* samples than in WT seedlings, suggesting profound changes in amino acid metabolism in response to a lowered ATP synthase abundance and activity in plants. The most dramatic differences were seen in the concentrations of glycine (Gly), ornithine, serine (Ser) and GABA. Grouping amino acids in families based on their synthesis pathways (Figure 7) and adding together their absolute abundances (Supplemental Table 6) showed that the aspartate family and aromatic amino acid family increased in abundance the least, pyruvate and glutamate family amino acids doubled in abundance, while Ser family amino acids increased 3 to 4-fold in RPF2-*atp1* compared to WT. On a percentage of the total amino acid pool basis this represented relative homeostasis for glutamate, aromatic and pyruvate family groups, but a one-third decrease for aspartate family and a 2 to 3-fold increase in Ser family amino acids. The latter represents 20-30% of the total amino acid pool as Ser family amino acids in RPF2-*atp1* compared to only 13% in WT. Notably, the Gly/Ser ratio increased nearly 4-fold in the RPF2-*atp1* mutants (Supplemental Table 7). Detailed analysis of the set of transcripts for *Arabidopsis* amino acid synthesis and degradation pathways (Hildebrandt, 2018) showed an induction of transcripts for tryptophan synthesis and degradation enzymes, but no consistent change in gene expression for enzymes in other amino acid metabolism families (Supplemental Table 8).

**Figure 7.**
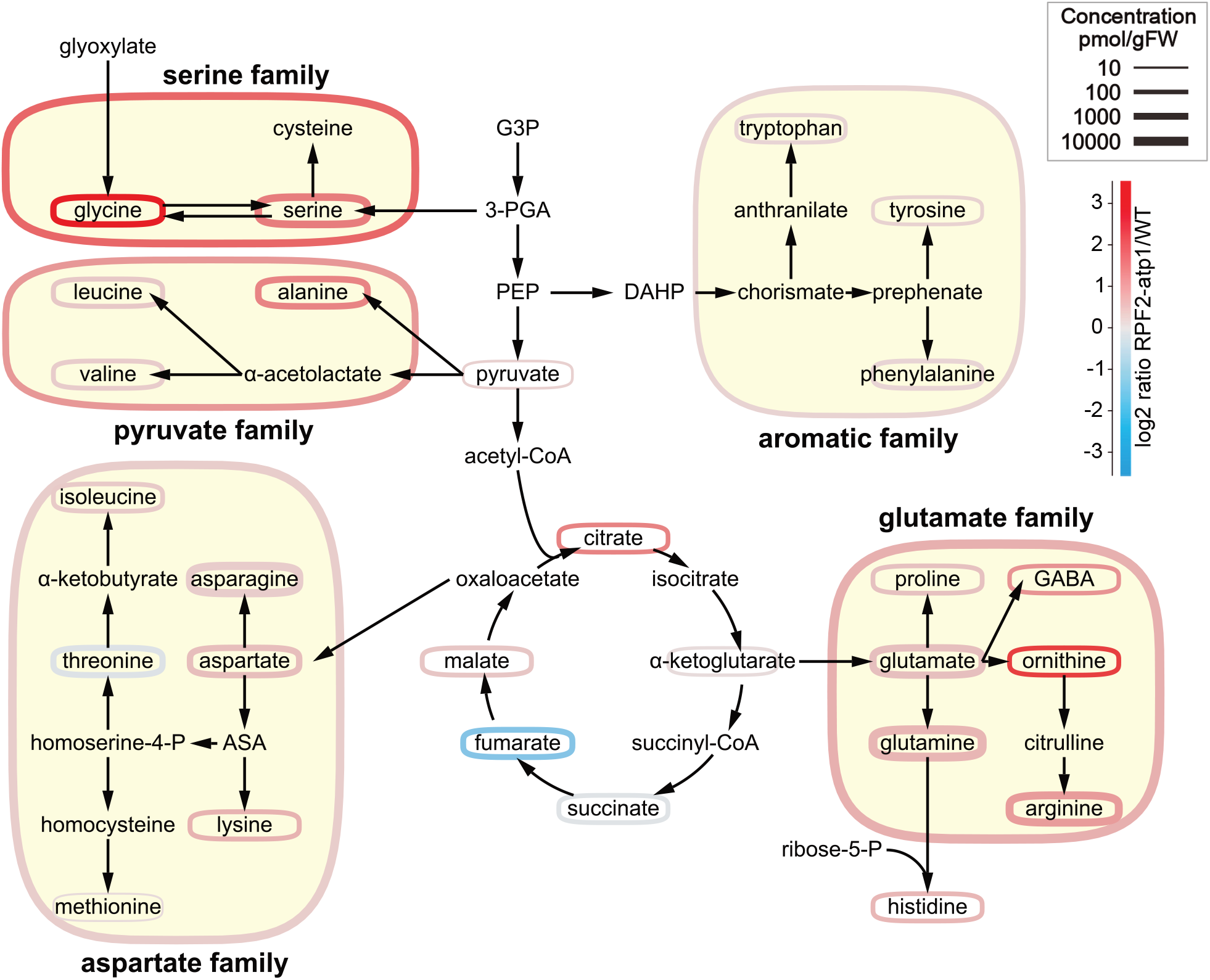
Perturbed Metabolite Abundances is in the RPF2-*atp1* Transformants. Simplified scheme of amino acid synthesis and TCA cycle, adapted from (Trovato et al., 2021). The amino acids are grouped by families according to their synthesis pathways. Colours represent the log2 ratio of RPF2-*atp1* to WT concentrations (see scale top right). Outline thickness is proportional to the log of the metabolite or metabolite pool concentration. ASA, aspartate semialdehyde; DAHP, 3-deoxy-D-arabinoheptulosonate-7-phosphate; G3P, glyceraldehyde 3-phosphate; 3-PGA, 3-phosphoglycerate; PEP, phosphoenolpyruvate.

## DISCUSSION

### Knockdown of Mitochondrial ATP Synthase Subunits

We succeeded in knocking down the mitochondrial ATP synthase by PPR-mediated cleavage of the *atp1* transcript, which led to levels of the Atp1 subunit that were 15-20% of WT (Figure 6B), compromising the assembly of the ATP synthase complex (Figures 1 and 3). Other reports show that a complete inactivation of mitochondrial ATP synthase during germination leads to seedling lethality due to insufficient ATP production during this energy-demanding process (Robison et al., 2009), prompting the choice for inducible and/or tissue-specific promoters to control the expression of the transgenes. Lethality has also been reported for mutants in the assembly factors of mitochondrial (ATP11 and ATP12) and plastid (ATP11, dual targeted) ATP synthases (Duan et al., 2020). Although we used the constitutive promoter NOS, we obtained a range of phenotypes, relating to differing levels of expression of the RPF2-*atp1* construct (Supplemental Figure 2), the most drastic being lethality. The phenotypic consequences of this inactivation in the RPF2-*atp1* plants were consistent with those reported for plants altered in the expression of nuclear subunits of the F_1_F_o_ ATP synthase such as slow growth, downward curled rosette leaf morphology, and male sterility (Robison et al., 2009; Geisler et al., 2012; Shaya et al., 2012; Li et al., 2019). We show that the assembled F_1_F_o_ ATP synthase in these plants was substantially lower in abundance than WT by western blot using a monoclonal antibody raised against Atp1 (single epitope) but the subunits could be accurately quantified as being 15-25% for both Atp1 and most subunits of the ATP synthase complex by quantitative mass spectrometry (Figure 6). These changes occurred with negligible variation in transcript abundance for any subunits other than Atp1 (Supplemental Figure 9).

### Independence of ATP synthase abundance from that of electron transport chain complexes

Our proteomics analysis, supported by BN PAGE and western blots, revealed that only the assembly and function of the ATP synthase complex was affected in the RPF2-*atp1* plants, as subunits of respiratory complexes I-IV were not significantly affected (Supplemental Figure 9, Supplemental Table 2). This observation was also recently reported for RNAi lines of the d subunit (Liu et al., 2021), suggesting that the biogenesis of the plant ATP synthase is independent from that of the other OXPHOS complexes. In that study, the authors observed a decrease in all ATPase subunits, except e (At3g01130, At5g15320), g (At4g29480, At2g19680, At4g26210) as we did, but also a (Atp6), *ε* (At1g51650) and ATP23 (At1g51680). In the present study, *ε* was down-regulated in the RPF2-*atp1* plants, like other structural components of the complex but we do not know if subunit a (Atp6) followed the same trend because it was not detected. The e and g subunits are thought to be recruited after an ATP synthase monomer is fully assembled (Röhricht et al., 2021), but as much less complex is formed in RPF2-*atp1*-9 and −16 mitochondria, it is possible that these 2 subunits can persist in the membrane in a residual F_o_ sub-complex. In *S. cerevisiae*, the assembly factor ATP23, a metallopeptidase located in the mitochondrial inner membrane, is involved in the processing of the a (Atp6) subunit and its assembly into the F_o_ module (Osman et al., 2007). *Arabidopsis* ATP23 was reported to have lost the ability to cleave the N-terminal extension of Atp6, but still acts as a chaperone to integrate Atp6 into the F_o_ module (Migdal et al., 2017). ATP23 (At3g03420) was not retained in our proteomics analysis after quality filtering, and its transcripts were not accumulated (Supplemental Table 2). Many CMS-related chimeric ORFs contain parts of genes encoding mitochondrial ATP synthase subunits, while others are adjacent to normal ATP synthase subunit genes (Hanson and Bentolila, 2004), supporting the energy deficiency hypothesis to explain the mechanism of CMS (Chen and Liu, 2014; Chen et al., 2017). However, how mitochondrial ATP synthase dysfunction leads to abnormal male gamete development remains unclear. As detailed in Table 1, the ATP synthase comprises nuclear as well as mitochondrially encoded subunits (Chen et al., 2017). So far, most efforts have been devoted to study the inactivation of nuclear ATP synthase subunit genes (Robison et al., 2009; Li et al., 2010; Geisler et al., 2012; Shaya et al., 2012). Antisense expression of ATP3 and ATP5 altered leaf morphology but not fertility, possibly because the required threshold of expression reduction might not have been reached (Robison et al., 2009). Mutation in the F_A_d subunit gene caused the lethality of pollen grains at a late developmental stage and the destruction of mitochondria in pollen grains (Li et al., 2010), consistent with the fact that ATP synthase dimers are the basis of the mitochondrial inner membrane morphology (Gu et al., 2019).

### Links between ATP synthase, mitochondrial dysfunction and retrograde regulation

Our data provides evidence linking a lowered mitochondrial ATP synthesis rate with general stress responses including induction of *AOX1a* and *AOX1d* gene expression and AOX abundance (Figure 3B, Supplemental Figure 9) and a degree of reduced fertility. This suggests that, as was observed in previous studies (Busi et al., 2011; Geisler et al., 2012; Liu et al., 2021), the constitutive reduction of ATP synthase activity by PPR-mediated knock-down of the *atp1* gene can trigger mitochondrial retrograde regulation (Van Aken et al., 2009; Schwarzlander et al., 2012; De Clercq et al., 2013). Our analysis suggests that two partly independent transcriptional pathways are activated in the *atp1* mutants (Figure 4): one is the typical ANAC017-dependent mitochondrial stress pathway, the other involves transcripts that are on the contrary, repressed by ANAC017 (also repressed by SNRK1α). This study also reports down regulation of male or female gametophyte-specific genes (Figure 4).

Whilst the exact signalling components linking loss of function of mitochondrial ATP synthase to primary metabolism and development during vegetative and reproductive stages remain to be identified (Meng et al., 2019), the retrograde signalling pathways of energy, hormone and stress responses activated in the RPF2-*atp1* plants are interesting research targets (Ng et al., 2014; de Souza et al., 2017).

Plants are very adaptable and use alternative pathways to compensate for deficiencies and maintain homeostasis. Despite having lower amounts of functional mitochondrial ATP synthase and substantially reduced rates of mitochondrial ATP synthesis, the RPF2-*atp1* plants maintain near normal concentrations of ATP in vegetative tissues, as also observed for δ subunit RNAi plants (Geisler et al., 2012). On the other hand, levels of ATP and ADP were reported to be decreased by 35% in the flowers of *atp9* inactivated *Arabidopsis* plants (Busi et al., 2011), but the ATP/ADP ratio was maintained. The authors suggest that the slow growth of the plants is not due to an energy deficit, but to the long-term metabolic effect of trying to maintain energy homeostasis, and they attribute the male and female fertility defects to the difficulty to maintain that homeostasis in tissues with high energy demand such as reproductive tissues.

### Links between lowering ATP synthase and leaf metabolism

Knockdown of expression of the gene encoding the delta subunit by RNA interference resulted in partially assembled ATP synthase and defects in pollen development coupled with broad relative changes in the abundance of metabolites (Geisler et al., 2012). We observed broadly similar organic acid and amino acid changes in our RPF2-*atp1* plants but by measuring absolute abundance (Supplemental Table 6) rather than relative abundance of amino acids (Geisler et al., 2012). We were also able to combine amino acids and changes in their abundance into amino acid synthesis families to show that the major effect is in the Ser family. The change represents not only a higher abundance but also a marked change in percentage of the Ser group to amino acids in other AA synthesis families and a four-fold increase in the Gly/Ser ratio (Figure 7, Supplemental Table 7). The conversion of Gly to Ser is a major source of NADH in leaf mitochondria and puts high demands on the respiratory chain as an electron sink (Kr<u>ömer</u>, 1995) and as a consequence is a primary source of energy for mitochondrial ATP generation in leaves. Gly accumulation in RPF2-*atp1* lines is thus most likely due to an acute slowing of Gly decarboxylation in mitochondria. Interestingly, the level of Gly accumulation in RPF2-*atp1*-9 was significantly higher than in RPF2-*atp1*-16 (Supplemental Table 6), consistent with the larger decrease in ATP synthase in RPF2-*atp1*-9 than in RPF2-*atp1*-16. Similar photorespiratory Gly decarboxylation limitations are also noted in mutants of AOX1a (Giraud et al., 2008), uncoupling protein UCP1 (Sweetlove et al., 2006) and NAD-MDH (Tomaz et al., 2010) suggesting that the loss of non-phosphorylating bypasses and other reductant dissipating pathways also limit photorespiration and lead to Gly accumulation in *Arabidopsis*. The upregulation of *AOX1a*, *AOX1d*, *NDA1*, *NDB2*, *NDB4* in RPF2-*atp1* could help to dissipate high levels of reducing equivalents; but may not be sufficient to allow adequate Gly decarboxylase activity in RPF2-*atp1* lines. There was no evidence of changes in expression of genes for GDC subunits in RPF2-*atp1*. While there was a marked induction of the cytosolic *SHM5* (At4g13890), it is a root specific isoform and its role in serine hydroxymethyl transfer is uncertain (Nogues et al., 2022)(Supplemental Table 2).

### Methods for Targeted Knockdown of Mitochondrial Genes

Cytoplasmic male sterility has been widely used in hybrid seed production, and many CMS-related genes were found by comparing the mitochondrial genomes of sterile and fertile plants (Chen et al., 2017; Kim and Zhang, 2018). But few of the candidate CMS-related genes have been functionally validated because of the lack of mitochondrial transformation strategies (Kazama et al., 2019). In recent years, three methods were proposed to target knockdown mitochondrial transcripts or genes via synthetic ribozymes (Val et al., 2011; Sultan et al., 2016; Niazi et al., 2019), designed PPR proteins (Colas des Francs-Small et al., 2018), or mito-TALENs (Kazama et al., 2019). Although proposed first (Val et al., 2011), the synthetic ribozyme strategy has not been widely used because of the complex chimeric structure of tRNA-like and custom ribozymes. Nevertheless, the knockdown of MatR by this method allowed a better understanding of this elusive but essential maturase encoded in a mitochondrial intron (Sultan et al., 2016). The mito-TALEN method gave the first direct evidence that *orf79* in rice and *orf125* in *Brassica* are the causes of CMS (Kazama et al., 2019). The knock-down of *atp1* in mitochondria via the redesigned RPF2 resulted in delayed growth and partial sterility in *Arabidopsis*, due to the lack of α subunit, and therefore of functional ATP synthase, indicating a highly efficient knock-down effect of the target gene by the designed PPR protein. TALEN-based technologies provide a permanent total knock out via DNA alteration (Boettcher and McManus, 2015) but designed PPR proteins provide a potentially reversible approach to knockdown mitochondrial genes by targeting RNA (Colas des Francs-Small et al., 2018). Different levels of knockdown effect can be achieved in T1 selection by using the designed PPR protein (Figures 2 and 3). Those phenotypes can be transmitted through generations after T1 selection (Figure 1C), providing versatile materials for laboratory research or breeding applications (Supplemental Figures 3 and 5). The use of inducible or tissue specific promoters would help modulate the effects of the transgene further, which is essential to study vital functions such as ATP production.

### PPR Protein Design for Organelle Biotechnology

The PPR code (Barkan et al., 2012; Miranda et al., 2018; Yan et al., 2019; Bernath-Levin et al., 2021) describing sequence specific binding ability between PPR proteins and their organelle target RNAs makes custom design usable for organelle biotechnology (Colas des Francs-Small et al., 2018). We show here that an engineered PPR RFL protein can be used to cleave a new target transcript within the coding sequence of *atp1*. Recently, engineered PPR10 proteins in combination with their RNA targets were used to activate plastid transgenes (Rojas et al., 2019), resulting in a ~40-fold increase in accumulation of the foreign proteins. That research established a new method using chloroplasts as biofactories to synthesize and store valuable biological molecules. With more natural PPR proteins being functionally validated, the artificially designed PPR proteins have potential to expand their application in organelles (Bernath-Levin et al., 2021; Royan et al., 2021).

## METHODS

### Protein Designing, Gene Cloning and Transformation

The designed RPF2-*atp1* gene with a 3xFLAG tag in C-terminal, as well as a fragment containing the NOS promoter and the coding sequence of the 25 amino acid *Solanum tuberosum* formate dehydrogenase (FDH) mitochondrial targeting peptide (Colas des Francs-Small et al., 1993) were commercially synthesized, and cloned between the *Eco*RI and *Bam* HI sites of pCAMBIA1380 binary vector (Bevan, 1984) by Gibson assembly. The synthetic genes were transferred to *Agrobacterium tumefaciens* and introduced into *A. thaliana* plants by floral dip (Clough and Bent, 1998).

### Primary Mutant Screening

For primary mutant screening, thirty-two T1 RPF2-*atp1* plants were grown in chambers with wild type plants (WT), plants expressing native RPF2 and the complex-V-deficient editing mutant *opt87* (Hammani et al., 2011) as controls. Genomic PCR was performed with primers specific for the construct (FDHPre3F and RPF2 410R) (Supplemental Table 9).

### Total RNA extraction and Northern Blotting

Total RNA was extracted from 4-week-old rosette leaves using PureZol kits (Bio-Rad). Eight micrograms of total RNA were run on a 1.2% denaturing agarose gel and transferred onto Hybond N + membrane (Amersham). Northern blotting was performed as described previously (Baudry et al., 2022) using oligonucleotide probes labelled at the 5′ end with biotin (Supplemental Table 9). The membranes were prehybridized for 1–2 h at 50 °C in 5 × SSC, 7% SDS, 100 μg·ml^−1^ heparin, 20 mM Na_2_HPO_4_ (pH 7.5) and hybridized overnight in the same buffer containing 1 nM biotinylated probe. Three short washes in 3 × SSC, 5% SDS, 25 mM Na_2_HPO4 pH 7.5 were performed at room temperature. The northern blots were developed with the Pierce Chemiluminescent Nucleic Acid Detection Module Kit (ThermoFisher Scientific).

### Mitochondrial RNA extraction and Circular RT-PCR

Crude preparations of mitochondria were isolated from 4-week-old Col-0 and RPF2-*atp1* seedlings grown on half-strength Murashige and Skoog medium as previously described (Colas des Francs-Small et al., 2012). Mitochondrial RNA was purified from the mitochondrial pellets using PureZol (Bio-Rad) and for each reaction, 2–3 μg were treated with Turbo DNase (Ambion). RNA was circularized using T4 RNA ligase and reverse transcription was performed with the Superscript III (Invitrogen) using specific primers (*atp1* RT-407R for the full transcript or the fragment upstream of the cleavage site and *atp1* RT5BR for the fragment downstream of it). PCR was performed with a nested primer (*atp1* cRT-195R or cRT-5DR) and a specific forward primer (*atp1* cRT-2F or *cRT-3F*) (Supplemental Table 9). PCR products were sequenced by Macrogen and the sequences aligned in Geneious Prime 2020.0.4).

### RT-qPCR

Total RNA was isolated from 2-week-old seedlings grown on plates under 16-hour photoperiod using RNAzol reagent (Sigma-Aldricht, Merck) according to the manufacturer’s instructions and DNA precipitation with a 4-fold volume of bromoanisole. Reverse transcription was performed on 1 microgram of RNA using random hexamers as previously described (Baudry et al., 2022).

The primers used are detailed in Supplemental Table 9.

### Protein Electrophoresis and Western Blotting

BN-PAGE, SDS-PAGE and western blotting were performed as previously described (Colas des Francs-Small et al., 2014; Vincis Pereira Sanglard and Colas des Francs-Small, 2022). The antibodies used in this work are listed in Supplemental Table 10.

### RNA Sequencing

Total RNA was isolated from 6-week-old rosette leaves and bolting flower buds of Col-0 and one of the RPF2-*atp1* (RPF2-*atp1*-9) transformed lines (T3) with PureZol reagent (BioRad). Three independent libraries for each genotype were made from total RNA treated with 250 ng of Turbo DNase (Ambion) using an Illumina TruSeq Stranded library preparation kit with Ribo-zero plant and random-primed reverse transcriptase. Sequencing (150bp, paired ends) was performed on an Illumina HiSeq4000 sequencer by Novogene (Novogene.com). RNA-seq data analysis methods are described in Supplemental Experimental Procedures.

### Proteomics Analysis

For quantitative untargeted mass spectrometry, crude mitochondrial pellets were obtained from 3-week-old WT, RPF2 native, RPF2-*atp1-9* and RPF2-*atp1-16* seedlings grown on plates and the samples from 3 independent experiments were prepared as previously described (Colas des Francs-Small et al., 2014; Petereit et al., 2020). Samples were analysed by LC-MS on a Thermo Exploris 480 mass spectrometer using data-dependent acquisition (see Supplemental Experimental Procedures).

### Adenylate Measurements

Absolute quantitation of AMP, ADP and ATP by LC-MS was carried out according to (Straube et al., 2021) with slight modifications (see Supplemental Experimental Procedures).

For the ATP synthesis rate, 20 μg of purified mitochondria isolated from 3-week-old native RPF2, RPF2 *atp1*-9 and RPF2 *atp1*-16 seedlings were equilibrated in 200 μl of respiration buffer (Meyer et al., 2009) containing 2 mM ADP. The respiration reactions (triplicates) were started by addition of NADH (1 mM final concentration) and stopped after 5 min by addition of 15% TCA. The adenylates were subsequently quantified according to the method described above. These samples were diluted 1/20 and the injection volume was 1 μL.

#### Mass Spectrometry Analyses of Organic Acids and Amino Acids

Four-week-old seedlings (~25 mg) grown under a 16-h light/8-h dark photoperiod were collected at the end of the night and snap-frozen in liquid nitrogen. Metabolites were extracted as previously specified (Lee et al., 2021). For LC-MS analysis of organic acids, sample derivatization was carried out based on previously published method with modifications (Han et al., 2013). Samples were analysed by an Agilent 1100 HPLC system coupled to an Agilent 6430 Triple Quadrupole (QQQ) mass spectrometer equipped with an electrospray ion source as described previously (Lee et al., 2021). For amino acid quantification, dried samples were resuspended in 50 ml water and analysed as described in (Le et al., 2021).

## DATA AVAILABILITY

The RNA-seq data have been deposited in the BioProject database, under the accession numbers PRJNA768306 (WT) and PRJNA893436 (RPF2-*atp1*-9).

Quantitative untargeted mass spectrometry data have been deposited to the ProteomeXchange Consortium via the PRIDE (Perez-Riverol et al., 2022) partner repository with the dataset identifier PXD037659. Reviewer account details: Username, Password: plVE9vk7

## FUNDING

This work was funded by the Australian Research Council (CE140100008, FL140100179).

## ACKNOWLEDGMENTS

No conflict of interest declared.

We thank Ricarda Fenske and Jacob Petereit from the University of Western Australia for their help during initial proteomics runs.

## AUTHOR CONTRIBUTIONS

C.C.d.F.-S. and I.S. conceived and designed the experiments. F.Y., L.V.P.S., E.S., C.P.L., G.G.K.O. and C.C.d.F.-S. carried out the experiments. F.Y., C.C.d.F.-S., S.S., E.S., C.P.L., G.G.K.O., A.H.M. and I.S. analysed the data and prepared the figures. F.Y., C.C.d.F.-S., A.H.M. and I.S. wrote the manuscript with editing contributions from all authors.

